# Prominin-1-Radixin Axis controls hepatic gluconeogenesis by regulating PKA activity

**DOI:** 10.1101/785154

**Authors:** Hyun Lee, Dong-Min Yu, Jun Sub Park, Hwayeon Lee, Jun-Seok Kim, Seung-Hoi Koo, Jae-Seon Lee, Sungsoo Lee, Young-Gyu Ko

## Abstract

Prominin-1 (Prom1) is a major cell surface marker of cancer stem cells, but its physiological functions in the liver have not been elucidated. We analyzed the levels of mRNA transcripts in serum-starved primary *Prom1*^+/+^ and *Prom1*^-/-^ mouse hepatocytes using RNA-sequencing (RNA-seq) data, and found that CREB target genes were down-regulated. This initial observation led us to determine that the Prom1 deficiency inhibited cAMP response element binding protein (CREB) activation and gluconeogenesis, but not cyclic AMP (cAMP) accumulation, in glucagon-, epinephrine-, or forskolin-treated liver tissues and primary hepatocytes, and mitigated glucagon-induced hyperglycemia. Because Prom1 interacted with radixin, the Prom1 deficiency prevented radixin from localizing to the plasma membrane. Moreover, systemic adenoviral knockdown of radixin inhibited CREB activation and gluconeogenesis in glucagon-treated liver tissues and primary hepatocytes, and mitigated glucagon-elicited hyperglycemia. Based on these results, we conclude that Prom1 regulates hepatic PKA signaling via radixin functioning as an A kinase-anchored protein (AKAP).

## Introduction

In well understood glucagon-elicited signaling pathway glucagon binds to and stimulates glucagon receptor, a G-protein-coupled receptor, in the plasma membrane of hepatocytes. The stimulated glucagon receptor in turn activates G_sα_ and adenylyl cyclase (AC). The cyclic AMP (cAMP) produced by the activated AC binds to the regulatory domain of protein kinase A (PKA) to liberate the catalytic domain of PKA from the regulatory domain and activate the protein. Not only does activated PKA induces glycogenolysis by consecutively phosphorylating and activating phosphorylase kinase and glycogen phosphorylase b, but also stimulates gluconeogenesis by phosphorylating and activating cAMP response element binding protein (CREB). Subsequently, activated CREB induces the transcriptional activation of gluconeogenesis-related genes such as phosphoenolpyruvate carboxykinase (*Pck*) and glucose-6-phosphatase (*G6p*) (Altarejos & Montminy, 2011, Meinkoth, Alberts et al., 1993).

Detergent-resistant lipid rafts, which are composed of cholesterol and glycolipids and serve as a signaling center, facilitate the cascade of glucagon signaling by organizing glucagon receptor, G_sα_, AC, A kinase-anchored proteins (AKAPs), PKA and PKA substrates in close proximity (Delint-Ramirez, Willoughby et al., 2011, Head, Patel et al., 2006, Kim, Lee et al., 2010). AKAPs are scaffolding proteins that bind PKA and its substrates. Thus, AKAPs allow PKA and its substrates to coexist in the same place, such as the plasma membrane, mitochondria or nucleus, enabling effective signal transduction by cAMP (Dema, Perets et al., 2015, Wong & Scott, 2004).

The penta-transmembrane glycoprotein prominin-1, also called CD133, localizes in membrane protrusions such as microvilli, filopodia and primary cilia in epithelial cells, membrane expansions in the myelin sheath originating from oligodendrocytes, membrane invaginations in the outer segment of rod photoreceptor cells and the midbody in epithelial cells (Corbeil, Roper et al., 2001, Corbeil, Roper et al., 1999, Corbeil, Roper et al., 2000, Dubreuil, Marzesco et al., 2007, Zacchigna, Oh et al., 2009). Mice with a systemic deficiency in Prom1 exhibit disk dysmorphogenesis and photoreceptor degeneration along with a complete loss of vision, indicating that Prom1 might be necessary for the formation of membrane extrusions (Zacchigna et al., 2009). In addition, Prom1 has been extensively studied as a major cancer stem cell marker in human brain, colon, ovarian and liver tumors. The Prom1-positive cell population in these tumors has characteristics of self-renewal, differentiation potential and resistance to chemo- and/or radiotherapy as well as tumor development after xenograft transplantation in immunocompromised mice (Dalerba, Dylla et al., 2007, Krishnan, Ochoa-Alvarez et al., 2013, Li, Heidt et al., 2007). Because PI3K interacts with Prom1 and Akt is highly phosphorylated in Prom1-positive cell populations, the Prom1-PI3K-Akt signaling pathway may be required for the maintenance of cancer stem cells (Wei, Jiang et al., 2013). However, this pathway has not been verified in a Prom1-deficient animal model.

Although Prom1 is expressed at high levels in various epithelial cells in the brain, kidney, digestive track and liver, the physiological functions of Prom1 are poorly understood. We analyzed glucagon-elicited gluconeogenesis in the livers of Prom1-deficient mice to understand Prom1 function. Here, we report that Prom1 is required for glucagon-induced PKA activation and hyperglycemia through its interaction with radixin, which functions as an AKAP in the liver.

## Results

### Prom1 is required for glucagon- and cAMP-elicited gluconeogenesis in mouse primary hepatocytes

After confirming Prom1 expression in mouse liver as well as primary hepatocytes (Fig S1A), we analyzed the levels of mRNA transcripts in serum-starved primary *Prom1*^+/+^ and *Prom1*^-/-^ mouse hepatocytes using RNA-sequencing (RNA-seq) data to evaluate the transcriptional effects of Prom1 deficiency in the liver. We focused on the genes that were induced (*n* = 55) or repressed (*n* = 63) more than 2-fold with *p* < 0.05 (Fig 1A). Singular enrichment analysis of those 118 differentially expressed genes (DEG) for KEGG pathways revealed that pathways such as glycolysis/gluconeogenesis, glucagon and cAMP signaling pathway were highly enriched in the DEGs by Prom1 deficiency (Fig EV1B). Because the liver plays a central role in the regulation of glucose homeostasis, we focused on glycolysis/gluconeogenesis. Identified genes in these pathways included *G6pc* (glucose-6-phosphatase catalytic subunit), *Pck1* (phosphoenolpyruvate carboxykinase 1), *Pfkl* (phosphofructokinase), and *Pkm* (pyruvate kinase), and were consistently downregulated by Prom1 deficiency (Fig 1A). These observations were further confirmed by qRT-PCR (Fig 1B), immunoblotting (Fig 1C) and glucose uptake assays (Fig 1D), and the results led us to suspect that Prom1 may be involved in hepatic gluconeogenesis.

**Figure 1.**
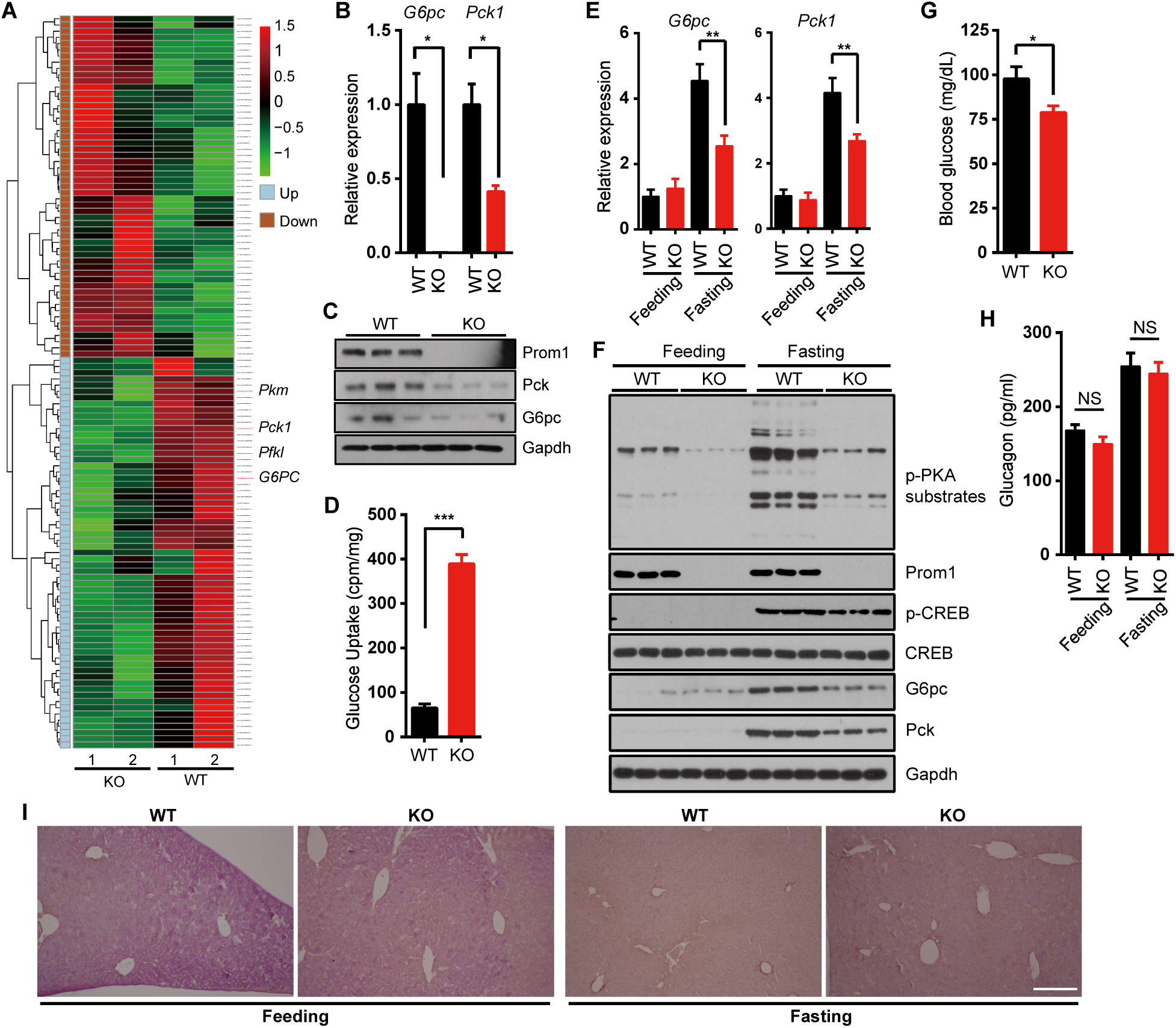
Prom1 deficiency prevents glucagon-induced gluconeogenesis. **A-D** Primary hepatocytes were isolated from 12-week-old male *Prom1*^+/+^ (WT) or *Prom1*^-/-^ (KO) mice, serum-starved and / or stimulated with glucagon. **A** Differentially expressed genes more than 2-fold with *p* < 0.05 (118 DEGs) induced (n=55) or repressed (n=63) by *Prom1* knockout (n = 2/group). *G6pc*, glucose-6-phosphatase catalytic subunit; *Pck1*, phosphoenolpyruvate carboxykinase 1. **B** Relative expression levels of *G6pc* and *Pck1* mRNAs were determined by RT-qPCR (n = 3/group) after serum starvation for 16 hours. \**p* < 0.05. **C** Levels of the Prom1, Pck, G6pc and Gapdh proteins were determined by immunoblotting. **D** The 8 hours serum-starved *Prom1*^+/+^ or *Prom1*^-/-^ hepatocytes were incubated with 2-[^3^H]deoxyglucose for 10 min. Glucose uptake was assessed by calculating the amount of cell-associated radioactivity normalized to the amount of protein. n = 3 mice per group. \*\*\**p* < 0.001. **E-I** Livers were isolated from fasted mice. **E** Levels of the *G6pc* and *Pck1* mRNAs in the liver of mice after 18 hours fasting were determined by quantitative PCR (qPCR). The level of each mRNA was normalized to the level of the *18s* rRNA. n = 3/fed, 7 or 8/fasted. \*\**p* < 0.01. **F** Levels of phospho-PKA substrates (p-PKA substrates), phospho-CREB (p-CREB), CREB, G6pc Pck, Prom1 and Gapdh were determined after 18 hours fasting. **G** Blood glucose levels (mg/dL) were measured after 24 hours fasting. n = 12 or 14/group. \**p* < 0.05. **H** Serum glucagon level in the fed or fasted mice was determined. n = 3/fed, 7 or 8/fasted. NS = not significant. **I** Glycogen contents in the liver of mouse in each condition was shown by PAS (periodic acid schiff) staining. Data information: Data are presented as mean values ± SEM. Two-tailed Student’s t-test was used for statistical analysis. Scale bar (**I**) = 500 μm.

To confirm the initial observations in mouse primary hepatocytes, we examined livers of *Prom1*^+/+^ and *Prom1*^-/-^ mice in the fasting state. Expression of *G6pc* and *Pck1* were decreased in *Prom1*^-/-^ mouse liver compared to the level in *Prom1*^+/+^ mouse liver in the fasting state (Fig 1E). Fasting-induced CREB phosphorylation was also decreased in *Prom1*^-/-^ mouse liver (Fig 1F). *Prom1*^-/-^ mouse had lower blood glucose level without difference in blood glucagon levels (Fig 1G and H). Hepatic glycogen breakdown during fasting was reduced in *Prom1*^-/-^ mice (Fig 1I). Livers of two groups did not show any histological difference (Fig S1C). These results suggested that loss of Prom1 interfered with the activation of hepatic gluconeogenesis.

To further confirm the involvement of Prom1 in hepatic gluconeogenesis, we determined the expression levels of hepatic gluconeogenic genes in *Prom1*^+/+^ and *Prom1*^-/-^mice hepatocytes after glucagon stimulation. The Prom1 deficiency interfered with the upregulation of *G6pc* and *Pck1* expression in glucagon-stimulated primary hepatocytes (Fig 2A and Fig S1D). Because glucagon induces the upregulation of gluconeogenic genes by activation of CREB, we measured glucagon-induced CREB phosphorylation. In Prom1-deficient hepatocyte, glucagon-induced CREB phosphorylation was significantly decreased (Fig 2B) and nuclear localization of CREB was also decreased as a result (Fig 2C and Fig S1E).

**Figure 2.**
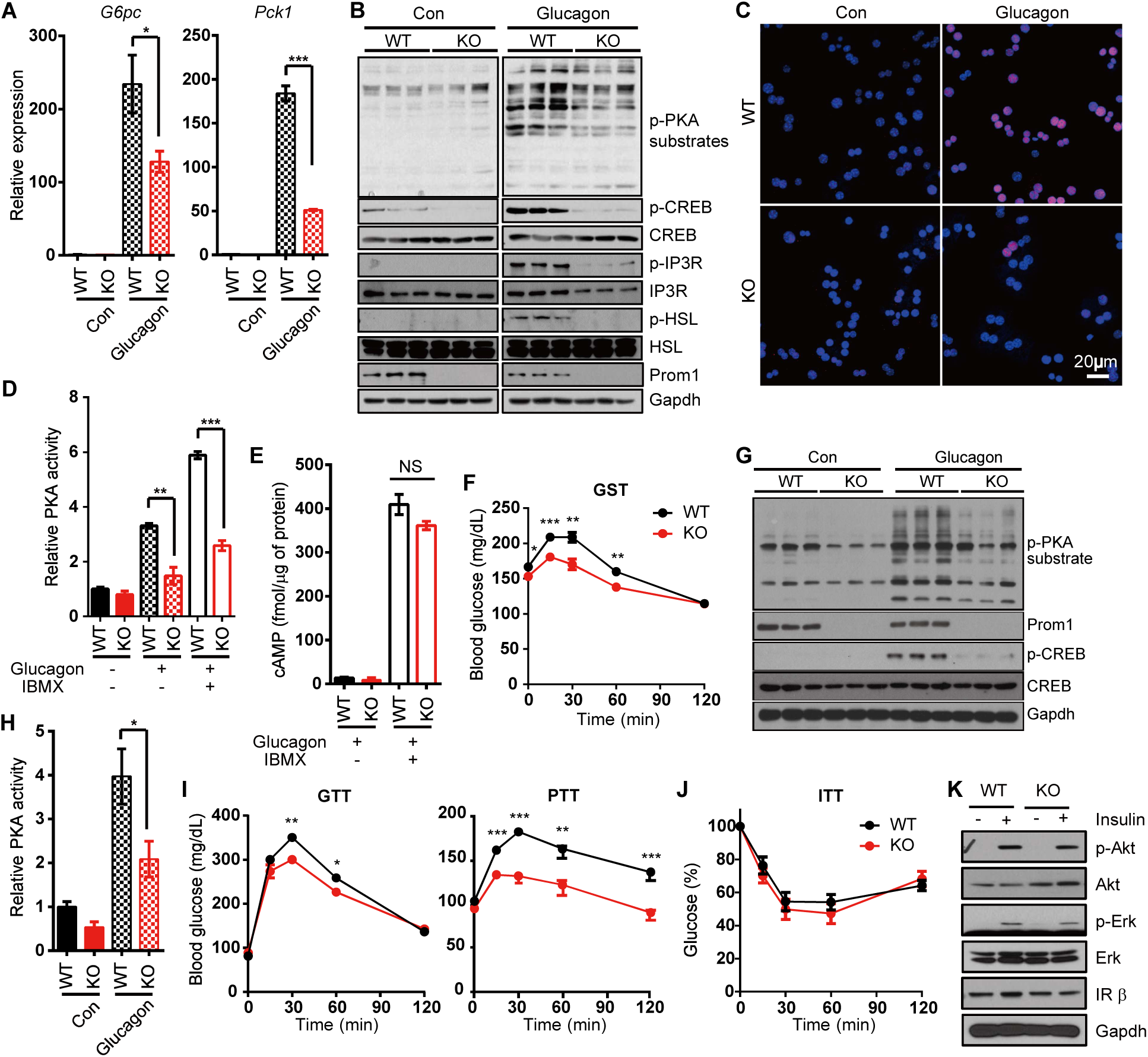
Prom1 deficiency prevents glucagon-induced hyperglycemia *in vivo*. **A-D** Primary hepatocytes were isolated from 12-week-old male *Prom1*^+/+^ (WT) or *Prom1*^-/-^ (KO) mice, serum-starved and stimulated with glucagon. **A** Levels of the *G6pc* and *Pck1* mRNAs were determined by quantitative PCR (qPCR) 2 hours after glucagon stimulation (10 nM). The level of each mRNA was normalized to the level of the *18S* rRNA. For these experiments, primary hepatocytes were cultured under 3D conditions using a Matrigel matrix. n = 3 mice per group. \**p* < 0.05; \*\*\**p* < 0.001. **B** Levels of phospho-PKA substrates (p-PKA substrates), phospho-CREB (p-CREB), CREB, phospho-Inositol trisphosphate receptor (p-IP3R), IP3R, phospho-Hormone-Sensitive Lipase (p-HSL), HSL, Prom1 and Gapdh were determined after a 10-min glucagon stimulation (10 nM). **C** The nuclear localization of p-CREB was determined by immunofluorescence staining after 10 min glucagon stimulation (10 nM). **D** Relative PKA activities after 10 min glucagon stimulation (10 nM) were determined using the PKA assay kit in the absence or presence of 10 μM IBMX. n = 3 mice per group. \*\**p* < 0.01; \*\*\**p* < 0.001. **E** The cAMP concentration after 10 min glucagon stimulation (10 nM) was determined using the cAMP assay kit in the absence or presence of 10 μM IBMX. n = 3 mice per group. NS = not significant **F-H** 12-week-old male *Prom1*^+/+^ (WT) and *Prom1*^-/-^ (KO) mice were fasted for 4 hours and intraperitoneally injected with glucagon. **F** Blood glucose levels (mg/dL) were measured 0, 15, 30, 60 and 120 min after glucagon stimulation (200 µg/kg body weight). n = 10/group. \**p* < 0.05; \*\**p* < 0.01; \*\*\**p* < 0.001. **G** Levels of p-PKA substrates, p-CREB, CREB and Gapdh in the liver were determined by immunoblotting after a 5 min glucagon treatment (2 mg/kg body weight). **H** Relative PKA activities in the liver were determined using the PKA assay kit after a 5 min glucagon treatment (2 mg/kg body weight). n = 3/group. \**p* < 0.05. **I, J** Glucose disposal rates were measured in 12-week-old male mice using glucose, pyruvate (**I**) and insulin (**J**) tolerance tests. n = 10 mice per group. \**p* < 0.05; \*\**p* < 0.01; \*\*\**p* < 0.001. **K** Levels of p-Akt, Akt, p-Erk, Erk, IRβ and Gapdh were determined by immunoblotting after insulin stimulation (10 nM). Data information: Data are presented as mean values ± SEM. Two-tailed Student’s t-test was used for statistical analysis.

The decreased CREB phosphorylation and the reduced upregulation of gluconeogenic genes in glucagon-induced *Prom1*^-/-^ mouse hepatocytes led us to examine the effect of the Prom1 deficiency on glucagon receptor signaling pathway. The Prom1 deficiency reduced glucagon-induced phosphorylation of PKA substrates (Fig 2B and D). The PKA activity and cAMP level in the presence or absence of IBMX (3-isobutyl-1-methylxanthine), a phosphodiesterase (PDE) inhibitor, were measured in glucagon-stimulated *Prom1*^+/+^ and *Prom1*^-/-^ primary hepatocytes to investigate the abilities of the AC and PDE enzymes required to produce and degrade cAMP, respectively. Blocking cAMP degradation by IBMX failed to restore PKA activity in *Prom1*^-/-^ mouse hepatocytes (Fig 2D). The Prom1 deficiency did not change glucagon-induced cAMP production or degradation (Fig 2E). We also examined the effect of Prom1 overexpression on glucagon-induced CREB phosphorylation and cAMP production in *Prom1*^-/-^ primary hepatocytes. Adenoviral Prom1 overexpression restored glucagon-induced CREB phosphorylation (Fig S1F), but did not affect glucagon-elicited cAMP production (Fig S1G).

To corroborate our observations, we tested two different PKA activators, forskolin (an AC activator) and 8-Br-cAMP (a non-degradable cAMP analogue) in *Prom1*^+/+^ and *Prom1*^-/-^ primary hepatocytes. Consistent with the previous results, the Prom1 deficiency prevented forskolin- and 8-Br-cAMP-induced CREB phosphorylation (Fig S2A and B) and reduced glucagon-induced phosphorylation of PKA substrates even in the presence of IBMX (Fig S2C) without changing glucagon-induced cAMP production (Fig S2D). Taken together, we concluded that Prom1 regulated cAMP-induced PKA activation in primary hepatocytes.

### Prom1 is necessary for glucagon-elicited hepatic gluconeogenesis *in vivo*

The requirement for Prom1 in glucagon-elicited gluconeogenesis was further analyzed *in vivo* by treating *Prom1*^+/+^ and *Prom1*^-/-^ mice with glucagon. We conducted glucagon challenge tests in 12-week-old mice. Glucagon-elicited hyperglycemia was mitigated in *Prom1*^-/-^ mice compared to that in *Prom1*^+/+^ mice (Fig 2F). The livers of Prom1-deficient mice also exhibited reduced glucagon-induced phosphorylation of CREB and PKA substrates (Fig 2G and H). However, Prom1-deficient mice did not display a change in glucagon-induced cAMP production in the liver (Fig S2E). Increased glucose internalization in *Prom1*^-/-^ mouse hepatocytes (Fig 1D) led us to hypothesize that Prom1-deficient mice would show decreased blood glucose level due to the reduced hepatic gluconeogenic capacity. To test this hypothesis, we performed glucose, pyruvate and insulin tolerance tests in *Prom1*^+/+^ and *Prom1*^-/-^ mice. Prom1-deficient mice displayed improved glucose and pyruvate tolerance (Fig 2I). Prom1 deficiency did not change insulin tolerance (Fig 2J), as insulin-induced signaling was not affected (Fig 2K). These results demonstrated that the Prom1deficiency prevented hepatic gluconeogenesis but did not affect insulin signaling.

### Prom1 regulates β-adrenergic receptor signaling *in vivo*

Because glucagon receptor is a member of the GPCR (G-protein coupled receptor) family, we hypothesized that Prom1 would also regulate other GPCR signaling that involves PKA activation. To test this hypothesis we investigated PKA signaling in *Prom1*^+/+^ and *Prom1*^-/-^ primary hepatocytes treated with isoprenaline, a β-adrenergic receptor agonist. The Prom1 deficiency prevented phosphorylation of CREB and PKA substrates (Fig 3A and B), but not cAMP production (Fig 3C) in isoprenaline-treated primary hepatocytes. To further validate that Prom1 regulates β-adrenergic receptor signaling, we examined blood glucose level and PKA activation in *Prom1*^+/+^ and *Prom1*^-/-^ mice after epinephrine injection. Epinephrine-induced hyperglycemia was mitigated (Fig 3D), and PKA activation was reduced (Fig 3E) in *Prom1*^-/-^ mice, when compared to that in *Prom1*^+/+^ mice. Because immobilization stress causes hyperglycemia via cholinergic muscarinic activation (Tajima, Endo et al., 1996), we performed immobilization test on *Prom1*^+/+^ and *Prom1*^-/-^ mice to test the effect of Prom1 deficiency on the signaling by endogenous epinephrine. We found that the immobilization stress-induced hyperglycemia and PKA activation were decreased in *Prom1*^-/-^, when compared to *Prom1*^+/+^ mice (Fig 3F and G). However, the reduced responses of *Prom1*^-/-^ mice to immobilization stress was not due to the different epinephrine levels in the serum of *Prom1*^+/+^ and *Prom1*^-/-^ mice (Fig 3H).

**Figure 3.**
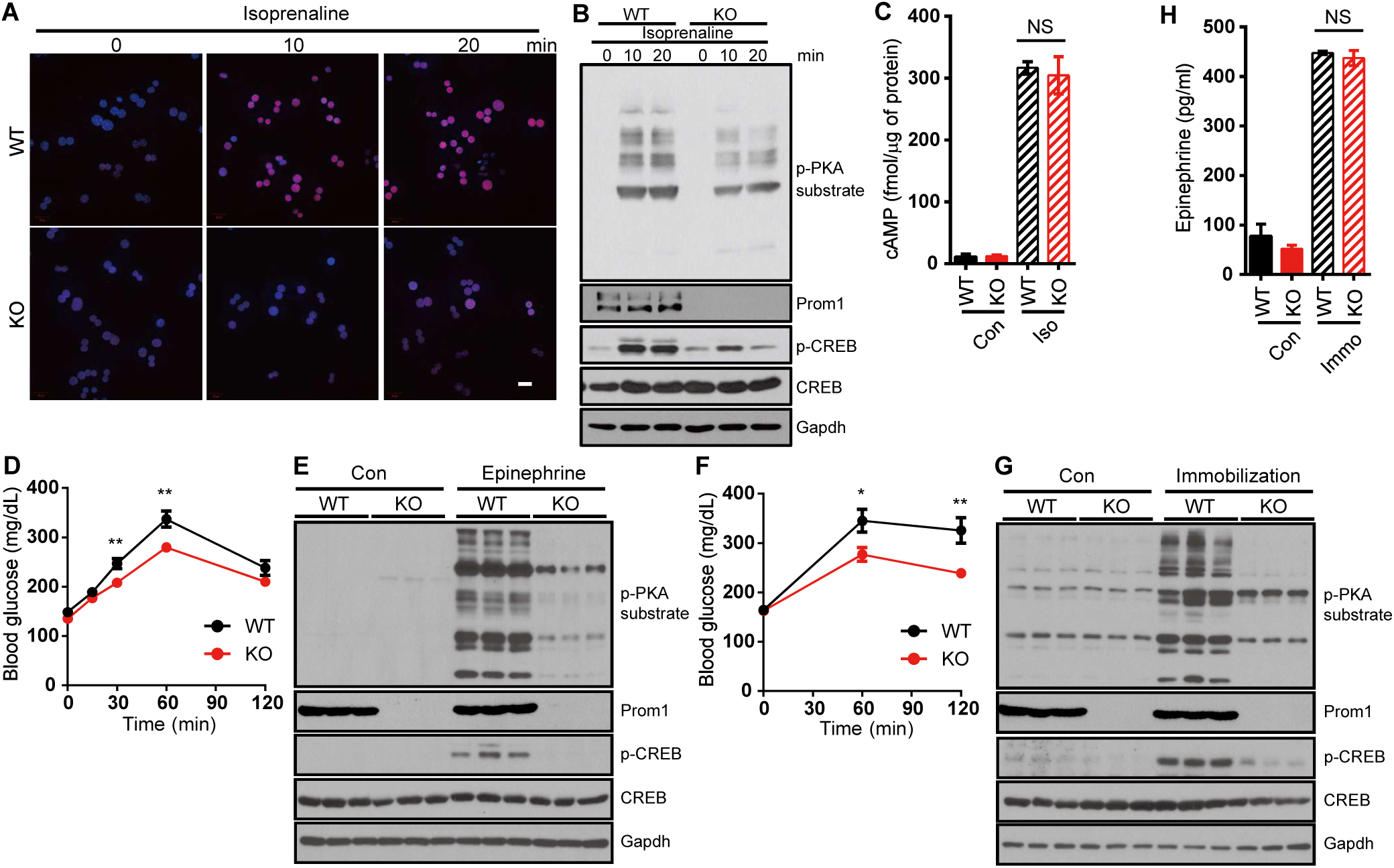
Prom1 deficiency prevents β-adrenergic receptor-mediated PKA activation. **A-C** Primary hepatocytes were isolated from 12-week-old male *Prom1*^+/+^ (WT) or *Prom1*^-/-^ (KO) mice, serum-starved for 16 h and stimulated with isoprenaline. **A** The nuclear localization of p-CREB was determined by immunofluorescence staining 10 min after isoprenaline stimulation (10 μM). **B** Levels of p-PKA substrates, p-CREB, CREB, Prom1 and Gapdh were determined by immunoblotting 0, 10 and 20 min after isoprenaline stimulation (10 μM). **C** The cAMP concentration was determined using the cAMP assay kit 10 min after isoprenaline stimulation (10 μM) in the presence of 10 μM IBMX. n = 3/group. NS = not significant. **D, E** *Prom1*^+/+^ (WT) and *Prom1*^-/-^ (KO) mice were fasted for 4 h and intraperitoneally injected with epinephrine. (3µg/10g). **D** Blood glucose levels (mg/dL) were measured 0, 15, 30, 60 and 120 min after epinephrine stimulation. n = 13 or 14/group. **p < 0.01. **E** Levels of p-PKA substrates, p-CREB, CREB and Gapdh in the liver were determined by immunoblotting after a 15 min epinephrine treatment. **F-H** *Prom1*^+/+^ (WT) and *Prom1*^-/-^ (KO) mice were fasted for 4 h and subjected to immobilization test. **F** Blood glucose levels (mg/dL) were measured 0, 60 and 120 min after immobilization. n = 10/group. *p < 0.05; **p < 0.01. **G** Levels of p-PKA substrates, p-CREB, CREB and Gapdh in the liver were determined by immunoblotting after a 30 min immobilization. **H** Serum epinephrine level of mice after 30min immobilization was determined. n = 3/control group, 7 or 8/immobilized group. NS = not significant. Data information: Data are presented as mean values ± SEM. Two-tailed Student’s t-test was used for statistical analysis.

### Radixin is the AKAP required for glucagon-induced PKA activation

Because Prom1 deficiency decreased PKA activation in response to glucagon without changing cAMP production (Figs 1 and 2), we postulated that Prom1 might regulate an AKAP which binds to the regulatory subunit of PKA and confines the PKA holoenzyme to a specific cellular location. We identified several AKAPs that are expressed in the liver using an RNA-seq analysis (Fig S3A). Among these AKAPs, AKAP7, 8, 8I, 9, 10, 12 and 13 were selected because they are cytoplasmic or plasma membrane-bound proteins that may interact with Prom1. In addition, Radixin was also selected as the AKAP in question among ERM (ezrin, radixin and moesin) protein family, because radixin is dominantly expressed in hepatocytes (Fig S3B), links actin to the plasma membrane in the liver (Kikuchi, Hata et al., 2002), and is known to function as an AKAP (Gloerich, Ponsioen et al., 2010, Hochbaum, Barila et al., 2011). We knocked down AKAPs expressed in the liver individually and examined their effects on glucagon-induced phosphorylation of PKA substrates in primary hepatocytes to identify which AKAP was regulated by Prom1. Glucagon-induced phosphorylation of PKA substrates was inhibited most markedly by small interfering RNAs (siRNAs) targeting Prom1 or radixin, suggesting that radixin might be an AKAP regulated by Prom1 (Fig 4A and Fig S3C and D). Because ezrin, a well-studied AKAP, was marginally expressed in the liver (Fig S3B), we tested the effect of ezrin knockdown by siRNA on glucagon-induced phosphorylation of PKA substrates as a control. Ezrin deficiency did not change the phosphorylation of PKA substrates (Fig S3E). Glucagon stimulation did not change the amount of radixin, PKA catalytic subunit or regulatory subunit (Fig S3F) as well. Similarly, adenoviral knockdown of radixin or Prom1 using short hairpin (sh) RNA in WT hepatocytes prevented glucagon-induced phosphorylation of CREB and other PKA substrates (Fig 4B, C, Fig S3G and Fig S4A). In contrast, radixin knockdown did not alter glucagon-induced cAMP production (Fig S4B). It has been reported that plasma membrane localization of Epac1, exchange factor directly activated by cAMP, and subsequent Rap1 activation is also mediated by radixin as an AKAP (Hochbaum et al., 2011). We determined Epac1 activity in Prom1^-/-^ primary hepatocytes treated with glucagon. Prom1 deficiency decreased Rap1 activation as well. (Fig S4C). These results suggested that radixin functions as the AKAP regulated by Prom1 in primary mouse hepatocytes.

**Figure 4.**
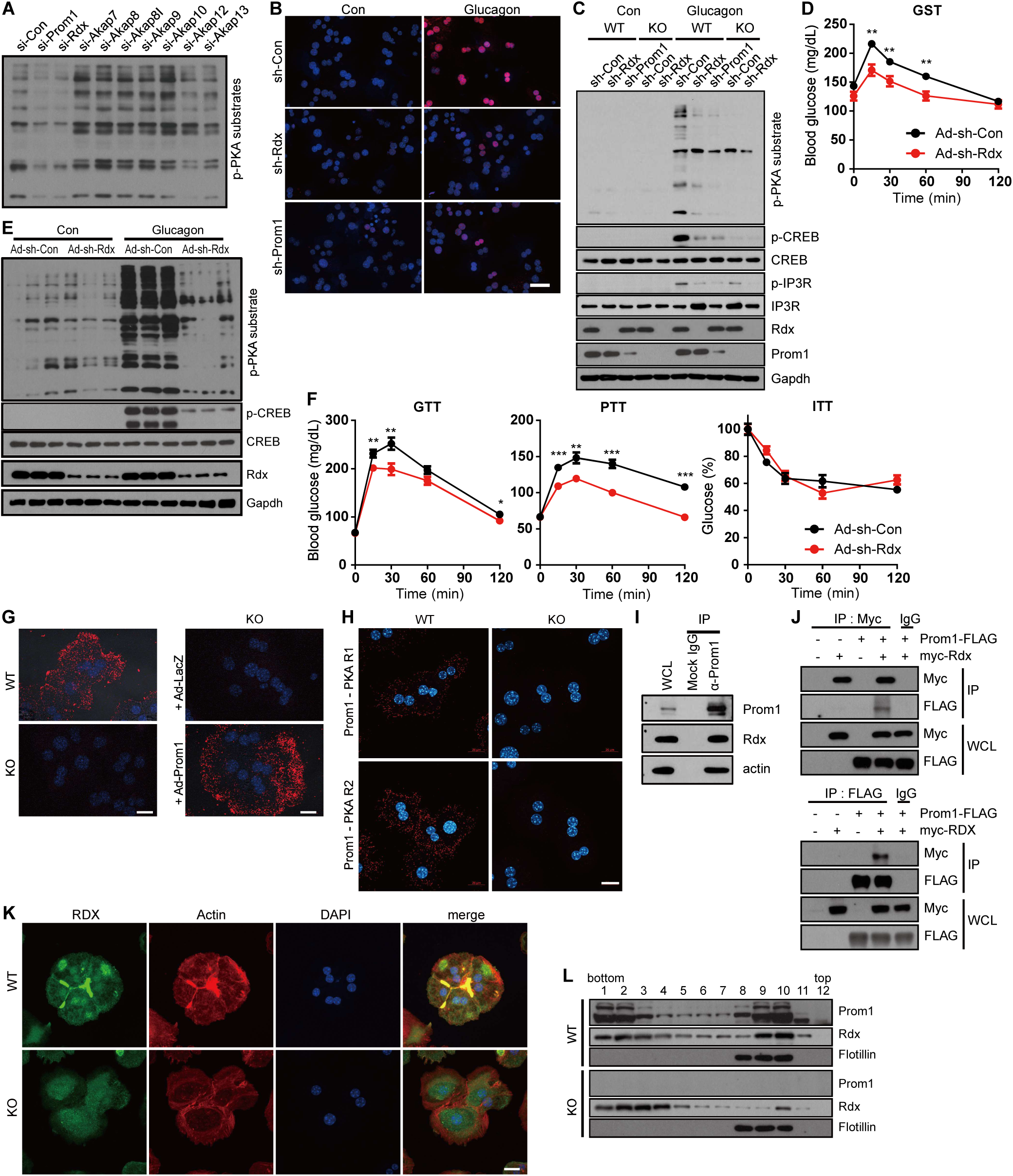
Radixin is the AKAP required for glucagon-induced PKA activation and hyperglycemia *in vivo* and Prom1 is required to confine radixin and actin. **A** The expression of Prom1, radixin, AKAP7, AKAP8, AKAP8I, AKAP9, AKAP10, AKAP12 or AKAP13 was silenced in WT hepatocytes using siRNAs. Hepatocytes were then serum-starved for 18 hours and stimulated with glucagon (10 nM) for 10 min. The phosphorylation of PKA substrates was determined by immunoblotting. **B, C** Prom1 and radixin expression were silenced in *Prom1*^+/+^ (WT) and *Prom1*^-/-^ hepatocytes (KO) by infection with an adenovirus harboring sh-control (sh-Con), sh-radixin (sh-Rdx) or sh-Prom1 for 24 h. Hepatocytes were further serum-starved for 18 h and stimulated with glucagon (10 nM) for 10 min. **B** The nuclear localization of p-CREB was determined by immunofluorescence staining after stimulation with glucagon. Blue; DAPI, Red; p-CREB. The percentage of cells with nuclear p-CREB staining among more than 300 cells in each group was also statistically analyzed (Supplementary Fig. 3g). **C** Levels of p-PKA substrates, p-CREB, CREB, p-IP3R, IP3R, Prom1, radixin and Gapdh were determined by immunoblotting after a 10-min glucagon stimulation. **D-F** 12-week-old male wild type mice were infected with an adenovirus harboring sh-control (sh-Con) or sh-radixin (sh-Rdx) for 3 days, fasted for 4 h and intraperitoneally injected with glucagon. **D** Blood glucose levels (mg/dL) were measured 0, 15, 30, 60 and 120 min after glucagon (200 µg/kg body weight) administration. n = 10 mice per group. \**p* < 0.05; \*\*\**p* < 0.001. **E** Levels of p-PKA, p-CREB, CREB, radixin and Gapdh in the liver were determined by immunoblotting after a 10-min glucagon (2 mg/kg body weight) stimulation for 10 min. **F** Glucose disposal rates in sh-Con or sh-Rdx (radixin) mice were measured using glucose, insulin and pyruvate tolerance tests. n = 10 mice per group. \**p* < 0.05; \*\**p* < 0.01; \*\*\**p* < 0.001. **G, H** Primary hepatocytes were obtained from 12-week-old male *Prom1*^+/+^ (WT) and *Prom1*^-/-^ (KO) mice or *Prom1*^-/-^ hepatocytes were infected with an adenovirus harboring LacZ (Ad-LacZ) or Prom1 (Ad-Prom1) for 24 h. The molecular interaction between Prom1 and radixin **G** or Prom1 and PKA regulatory subunits **H** in these hepatocytes was determined by a proximity ligation assay using anti-Prom1 and anti-radixin antibodies (**G**) or anti-Prom1 and anti-PKA regulatory subunits (**H**), respectively. **I** The molecular interaction between Prom1 and radixin in *Prom1*^+/+^ hepatocytes was determined by co-immunoprecipitation with an anti-Prom1 antibody or isotype control IgG. **J** HEK293 cells were transiently transfected with the combination of Prom1-Flag and Myc-radixin plasmids for 24 h. The molecular interaction between Prom1-Flag and Myc-radixin was determined by co-immunoprecipitation using isotype control IgG, anti-Flag or anti-Myc antibodies. **K** The cellular localization of radixin and actin in *Prom1*^+/+^ and *Prom1*^-/-^ hepatocytes was determined by immunofluorescence staining for radixin and phalloidin staining. **L** Detergent-resistant lipid rafts were isolated from *Prom1*^+/+^ and *Prom1*^-/-^ hepatocytes. Levels of the Prom1, radixin and flotillin proteins were determined in each fraction after sucrose gradient ultracentrifugation using immunoblotting. Data information: Data are presented as mean values ± SEM. Two-tailed Student’s t-test was used for statistical analysis. Scale bar (b) = 20 μm.

Next, to determine whether radixin functions as an AKAP during glucagon-induced gluconeogenesis *in vivo*, we knocked down radixin in mice using adenoviral shRNA, confirmed its transduction efficiency to be close to 100 % in hepatocytes (Fig S4D), and performed the glucagon challenge test. Glucagon-induced hyperglycemia was mitigated by radixin knockdown (Fig 4D). Moreover, radixin knockdown reduced the phosphorylation of CREB and PKA substrates (Fig 4E and Fig S4E), but did not alter cAMP production in the livers of glucagon-treated mice (Fig S4F). Glucose, pyruvate, and insulin tolerance tests were performed on radixin-knockdown mice to further investigate whether radixin regulates hepatic gluconeogenesis *in vivo*. Radixin knockdown improved glucose and pyruvate tolerance but not insulin tolerance (Fig. 4F), suggesting that radixin knockdown prevented glucagon-induced hepatic gluconeogenesis without affecting insulin signaling.

### Prom1 is necessary to confine radixin to the plasma membrane

To determine the mechanism by which Prom1 regulates the AKAP activity of radixin we first examined whether the two proteins were located in close proximity using a proximity ligation assay. Fluorescence signals were observed in *Prom1*^+/+^ but not in *Prom1*^-/-^ primary hepatocytes (Fig 4G, two left panels) and adenoviral overexpression of Prom1 drove the proximity-based fluorescence to reappear in Prom1^-/-^ primary hepatocytes (Fig 4G, two right panels). PKA regulatory subunits were also in close proximity to Prom1 (Fig 4H) implying the formation of Prom1/radixin/PKA complex. Endogenous Prom1 was co-immunoprecipitated with radixin in primary mouse hepatocytes (Fig 4I), suggesting the molecular interaction between Prom1 and radixin. Reciprocal immunoprecipitations of exogenously expressed Prom1 and radixin showed similar results in HEK293 cells (Fig 4J). The molecular interaction between Prom1 and radixin prompted us to speculate that Prom1 is required to confine radixin to the plasma membrane and to link actin to the plasma membrane. We determined the cellular localization of radixin and actin in *Prom1*^+/+^ and *Prom1*^-/-^ primary hepatocytes to test this hypothesis. Radixin and actin were clearly observed in cell-cell contact sites called the canalicular membranes in *Prom1*^+/+^ hepatocytes but not in *Prom1*^-/-^ hepatocytes (Fig 4K). Moreover, the Prom1 deficiency prevented cortical actin from localizing to canalicular membranes, suggesting that Prom1 was required to confine radixin and actin to the plasma membrane. Because Prom1 and ERM proteins are present in detergent-resistant lipid rafts, we speculated that radixin enrichment in detergent-resistant lipid rafts may depend on Prom1 expression. We examined the presence of radixin in detergent-resistant lipid rafts from *Prom1*^+/+^ and *Prom1*^-/-^ primary hepatocytes to test this hypothesis. Prom1-deficient cells exhibited a reduced amount of radixin in the detergent-resistant lipid rafts (Fig 4L). These results demonstrated that Prom1 is required to confine radixin to plasma membrane lipid rafts.

### The FERM domain of radixin is necessary for Prom1-dependent gluconeogenesis

To determine which domain in each protein is required for the interaction, we constructed various deletion mutants of both Prom1 and radixin (Fig S5A and B). Co-immunoprecipitation after transient expression of both genes in HEK293 cells showed that the carboxy-terminal tail of Prom1 (IC3) interacted with the FERM domain (1-310) of radixin, because the full-length protein and carboxy-terminal tail of Prom1 co-immunoprecipitated the full-length protein and FERM domain of radixin (Fig 5A and B). Competitive co-immunoprecipitation showed a gradual decrease in the molecular interaction between Prom1 and radixin in proportion to the increasing expression level of the FERM domain in HEK293 cells (Fig 5C), indicating the specific interaction between two proteins through FERM domain. Indeed, the canalicular localization of endogenous radixin disappeared in cells overexpressing the FERM domain (Fig 5D). Next, we monitored glucagon-induced PKA activation after adenoviral overexpression of the FERM domain in primary mouse hepatocytes. Overexpression of the FERM domain prevented glucagon-induced phosphorylation and nuclear localization of CREB (Fig 5E and Fig S5C). Overexpression of the FERM domain prevented glucagon-induced phosphorylation of PKA substrates (Fig 5F), but did not affect glucagon-elicited cAMP production (Fig S5D). We further analyzed the dominant negative effect of the FERM domain on glucagon-elicited gluconeogenesis *in vivo*. Systemic adenoviral overexpression of the FERM domain mitigated glucagon-elicited hyperglycemia (Fig 5G). In addition, overexpression of the FERM domain prevented the glucagon-induced phosphorylation of CREB and PKA substrates in the livers of glucagon-treated mice (Fig 5H), and improved glucose and pyruvate tolerance but not insulin tolerance (Fig. 5I). Taken together, we concluded that the FERM domain of radixin is necessary for the interaction with Prom1 and its confinement to lipid rafts to facilitate radixin’s function as an AKAP during hepatic gluconeogenesis.

**Figure 5.**
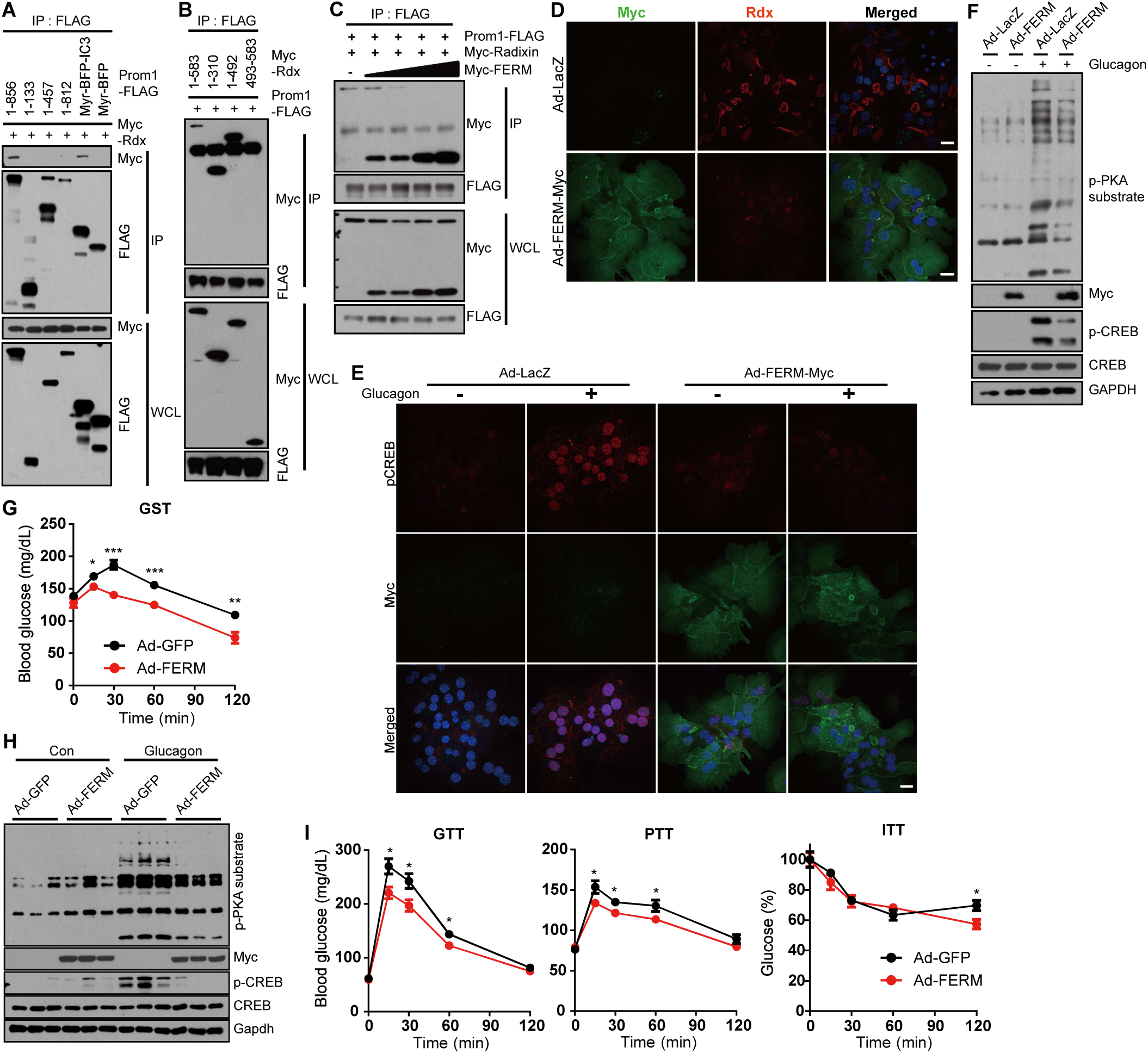
Overexpression of radixin mutant mitigates glucagon-elicited hyperglycemia. **A, B** The molecular interaction was determined by co-immunoprecipitation with an anti-FLAG antibody. HEK293 cells were co-transfected with plasmids expressing Myc-radixin and various deletion mutants of Prom1-Flag (**A**) or with plasmids expressing Prom1-Flag and various deletion mutants of Myc-radixin (**B**) for 24 h. **C** The molecular interaction between Prom1-Flag and Myc-radixin in the presence of increasing amounts of Myc-FERM was determined by co-immunoprecipitation with an anti-Flag antibody. **D-F** *Prom1*^+/+^ primary hepatocytes were infected with an adenovirus harboring FERM-Myc (Ad-FERM-Myc) or LacZ (Ad-LacZ) for 48 h. **D** The cellular localization of radixin was determined by immunofluorescence staining. **E** The nuclear localization of p-CREB was determined by immunofluorescence staining after glucagon stimulation for 10 min. **F** Ad-FERM-Myc- or Ad-LacZ-expressing *Prom1*^+/+^ hepatocytes were serum-starved for 16 h and stimulated with 10 nM glucagon for 10 min. Levels of p-PKA substrates, p-CREB, CREB, FERM-Myc and GAPDH were determined by immunoblotting. **G-I** Twelve-week-old male wild type mice were infected with an adenovirus harboring GFP or FERM for 3 days, fasted for 4 h and intraperitoneally injected with glucagon. **G** Blood glucose levels (mg/dL) were measured 0, 15, 30, 60 and 120 min after glucagon stimulation (0.2 mg/kg body weight). n = 10 mice per group. \**p* < 0.05; \*\**p* < 0.01; \*\*\**p* < 0.001. **H** Levels of p-CREB, CREB, p-PKA substrates, and FERM in the liver 10 min after glucagon stimulation (2 mg/kg body weight) were determined by immunoblotting. n = 3 mice per group. **I** Glucose disposal rates in GFP- or FERM-overexpressing mice were measured using glucose, pyruvate, and insulin tolerance tests. n = 10 per group. \**p* < 0.05; \*\**p* < 0.01. Data information: Data are presented as mean values ± SEM. Two-tailed Student’s t-test was used for statistical analysis. Scale bar = 20 μm.

AKAP activity of radixin in glucagon-elicited gluconeogenesis was further substantiated by knockdown-rescue experiment. We examined glucagon-induced PKA activation in primary mouse hepatocytes in which endogenous radixin was knocked down by adenoviral shRNA, and shRNA-resistant radixin^R^ or LPTD^R^ was re-introduced by adenoviral transduction. We used the LPTD mutant of radixin, because L421P mutation disrupts the ability of radixin to bind to PKA, therefore losing AKAP activity and T564D mutation is known to mimic phosphorylation in all ERM proteins in its open and active conformation (Deming, Campbell et al., 2015). Ectopic expression of shRNA-resistant radixin^R^ rescued glucagon-induced phosphorylation of PKA substrates, while radixin mutant LPTD^R^ could not (Fig 6A). Overexpression of LPTD^R^ mutant alone in primary mouse hepatocytes decreased the glucagon-induced phosphorylation of CREB and PKA substrates (Fig 6B), which suggested the dominant negative effect of LPTD^R^ mutant as well.

**Figure 6.**
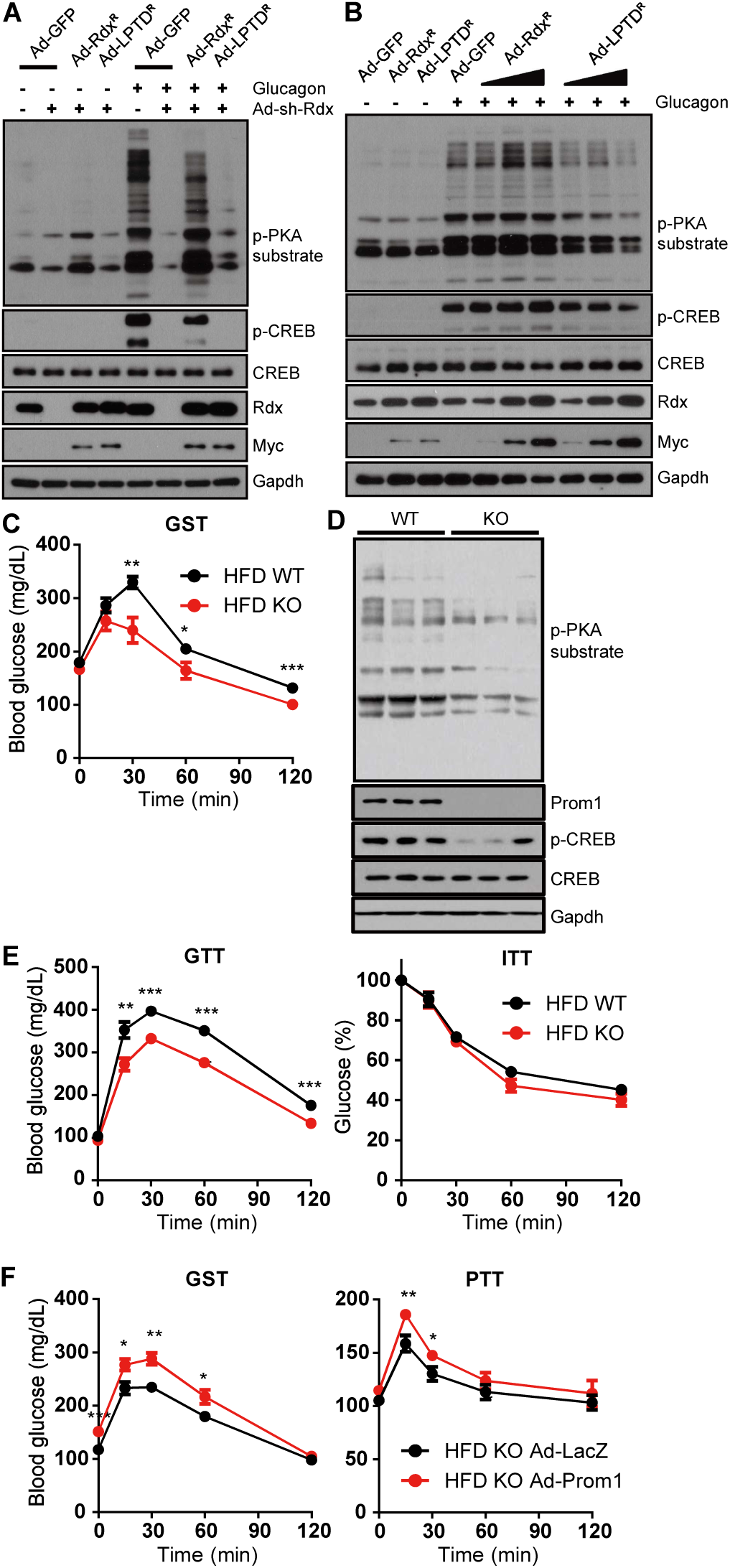
Prom1 disruption ameliorates glucagon-induced hyperglycemia in DIO mice. **A, B** *Prom1*^+/+^ primary hepatocytes were infected with an adenovirus harboring sh-control (sh-Con), sh-radixin (sh-Rdx), GFP (Ad-GFP), and/or sh-resistant wildtype radixin (Ad-Rdx^R^) or AKAP-dead LPTD mutant of radixin (Ad-LPTD^R^) for 24 h. Hepatocytes were further serum-starved for 18 hours and stimulated with glucagon (10 nM) for 10 min. Levels of p-PKA substrates, p-CREB, CREB, radixin, Myc-radixin and Gapdh were determined by immunoblotting **C-F** Four-week-old male *Prom1*^+/+^ and *Prom1*^-/-^ mice fed a high-fat diet for 8 weeks, were fasted for 4 h. **C** DIO *Prom1*^+/+^ and *Prom1*^-/-^ mice were intraperitoneally injected with glucagon. Blood glucose levels (mg/dL) were measured 0, 15, 30, 60 and 120 min after glucagon (100 µg/kg body weight) stimulation. n = 10 mice per group. \**p* < 0.05; \*\**p* < 0.01; \*\*\**p* < 0.001. **D** Levels of p-PKA substrates, p-CREB, CREB, Prom1 and Gapdh were determined by immunoblotting 10 min after glucagon (2 mg/kg body weight) stimulation. n = 3 mice per group. **E** Glucose disposal rates in DIO *Prom1*^+/+^ and *Prom1*^-/-^ mice were measured using glucose and insulin tolerance tests. n = 10 mice per group. \*\**p* < 0.01; \*\*\**p* < 0.001. **F** DIO *Prom1*^-/-^ mice were infected with an adenovirus harboring LacZ or Prom1 for 24 h, fasted for 4 h and intraperitoneally injected with glucagon (100 µg/kg body weight). Blood glucose levels (mg/dL) were measured 0, 15, 30, 60 and 120 min after glucagon stimulation (GST). Glucose disposal rates were measured using the pyruvate tolerance test (PTT). N = 10 mice per group. \**p* < 0.05; \*\**p* < 0.01; \*\*\**p* < 0.001. Data information: Data are presented as mean values ± SEM. Two-tailed Student’s t-test was used for statistical analysis

Because our results showed that Prom1-deficient mice displayed improved glucose and pyruvate tolerance but not insulin tolerance (Fig 2I and J), we examined glucagon-elicited gluconeogenesis in high-fat diet-induced obese (DIO) *Prom1*^+/+^ and *Prom1*^-/-^ mice. DIO *Prom1*^-/-^ mice exhibited a mitigation of glucagon-elicited hyperglycemia (Fig 6C). The Prom1 deficiency also prevented glucagon-induced phosphorylation of CREB and other PKA substrates (Fig 6D). In DIO *Prom1*^-/-^ mice, glucose tolerance, but not insulin tolerance, was improved (Fig 6E). Moreover, systemic adenoviral overexpression of Prom1 in DIO *Prom1*^-/-^ mice enhanced glucagon- and pyruvate-induced hyperglycemia (Fig 6F). These *in vivo* results showed that the Prom1 deficiency protected mice from diet-induced glucose intolerance

## Discussion

Although Prom1 is a cancer stem cell marker located in plasma membrane detergent-resistant lipid rafts, the cellular functions of Prom1 have remained elusive (Krishnan et al., 2013, Roper, Corbeil et al., 2000). For the first time, our study provides new insights into the physiological role of Prom1. We examined the mechanism by which Prom1 regulates hepatic gluconeogenesis using Prom1-deficient mice. The Prom1-radixin axis is a key signaling pathway that regulates cAMP-mediated PKA activation. We propose that Prom1 confines radixin to the plasma membrane and radixin recruits the PKA holoenzyme to the plasma membrane. Subsequently, glucagon-G protein-coupled receptor (GPCR)-G_sα_-AC signaling pathway produces cAMP, liberating and activating the PKA catalytic subunit from the PKA regulatory subunit attached to radixin. Because Prom1 deficiency interferes with the function of radixin as an AKAP, Prom1 deficiency prevents cAMP-mediated PKA activation.

ERM family proteins (ezrin, radixin and moesin) act as AKAPs because they interact with the PKA regulatory subunit and various plasma membrane-associated PKA substrates (Neisch & Fehon, 2011). Indeed, ERM proteins are required for PKA-mediated phosphorylation of different membrane-associated proteins, such as β-adrenergic receptor, cystic fibrosis transmembrane conductance regulator (CFTR), Na^+^-H^+^ exchanger 3 (NHE3), connexin 43 (Cx43) and exchange protein directly activated by cAMP (EPAC) (Bretscher, Edwards et al., 2002, Fouassier, Duan et al., 2001, Hochbaum et al., 2011, Pidoux, Gerbaud et al., 2014, Sun, Hug et al., 2000, Weinman, Steplock et al., 2003). We focused on the Prom1-radixin complex because among these ERM proteins radixin is the dominant ERM protein in hepatocytes (Kikuchi et al., 2002, Tsukita, Hieda et al., 1989). Radixin knockout mice show hyperbilirubinemia due to loss of multidrug resistance protein 2 (MRP2) from canalicular membranes in the liver (Kikuchi et al., 2002), deafness associated with progressive degeneration of cochlear stereocilia in the inner ear (Kitajiri, Fukumoto et al., 2004), and impaired reversal learning and short-term memory by modulating inhibitory synapse transmission (Loebrich, Bähring et al., 2006). Our study provides the first evidence to show that radixin is also involved in glucagon/β-adrenergic receptor-mediated gluconeogenesis via Prom1-radixin axis. Prom1 deficiency disrupts the localization of radixin in canalicular membranes, prevents the formation of cortical actin (Fig 4K) and inhibits glucagon-, isoprenaline- or cAMP-induced PKA activation (Fig 2, Fig 3, and Fig S2) without changing the amount of PKA catalytic subunit (Fig S3F). These results demonstrate that Prom1 deficiency abrogates the function of radixin as an AKAP which allows the PKA holoenzyme and PKA substrates to assemble in the same place. Failing to confine the PKA holoenzyme and PKA substrates in close proximity due to Prom1 deficiency may explain the reduced phosphorylation level of PKA substrates even at the supra-physiological concentration of activators used in our study. In addition, because PKA has generous substrate specificity (Dalerba et al., 2007), it is crucial to tightly regulate PKA localization through its interaction with AKAPs (Beene & Scott, 2007) and activation through confined cAMP concentration (Rich, Fagan et al., 2001, Zaccolo & Pozzan, 2002). Recently, Smith *et al*. (Smith, Esseltine et al., 2017) have demonstrated that local PKA activation is regulated by the conformational change of holoenzyme, not by the complete dissociation of catalytic subunit at the physiological concentration of cAMP. In accordance with these reports our findings also demonstrated that local confinement of PKA to which cAMP is readily available by the interaction with radixin, hepatocyte specific AKAP, and Prom1 complex is an important process that provides the specificity in the regulation of glucagon-induced gluconeogenesis. This may explain the lack of action from other AKAP proteins in the presence of glucagon or 8-Br-cAMP. More interestingly, we demonstrated that the importance of Prom1-radixin axis not only in glucagon-elicited gluconeogenesis, but also in β-adrenergic receptor-mediated gluconeogenesis (Fig. 3). These results suggest that Prom1-AKAP interaction may participate in regulating GPCR signaling in other phenotypes or organs.

Membrane-associated AKAPs are known to be involved in cAMP-induced phosphorylation and the activation of nuclear CREB. Although membrane-bound AKAP5 interacts with E-cadherin, β-adrenergic receptor, adenylyl cyclase and the cytoskeleton, cAMP-induced phosphorylation of nuclear CREB is increased by its overexpression but is reduced by its knockdown (Altier, Dubel et al., 2002, Fraser, Cong et al., 2000, Gorski, Gomez et al., 2005). The cAMP-induced phosphorylation of nuclear CREB is also reduced with a cell-permeable peptide treatment that inhibits the molecular interaction between PKA and AKAP (Friedrich, Aramuni et al., 2010, Godbole, Lyga et al., 2017). Similar to the observations in the above examples, the Prom1-radixin complex in the plasma membrane regulates cAMP-induced phosphorylation of nuclear CREB. However, the molecular mechanism by which radixin regulates the phosphorylation of nuclear CREB still remains elusive.

Using Prom1-deficient or radixin-knockdown mice, we show that the Prom1-radixin complex is required for hepatic gluconeogenesis. Because both lean and high-fat diet-fed mice lacking Prom1 or radixin exhibited a decreased blood glucose level after a 4-h fast and improved glucose and pyruvate tolerance without changing insulin sensitivity, Prom1 and radixin may be excellent target proteins for lowering the blood glucose level in patients with diabetes. Therefore, a chemical drug that interferes with the molecular interaction between Prom1 and radixin might thus be useful as a treatment for hyperglycemia by inhibiting gluconeogenic signaling pathway.

## Materials and Methods

### Animal studies

Prom1 knockout mice were purchased from The Jackson Laboratory (Stock NO. 017743, Bar Harbor, ME, USA). The *Prom1*^-/-^ mice were backcrossed with C57BL/6N mice for five generations. All animal studies were conducted with the approval of the Korea University Institutional Animal Care and Use Committee and the Korean Animal Protection Law (KUIACUC-2017-14 and -2018-6).

For glucagon stimulation tests, glucagon (200 μg/kg body weight for mice fed a normal chow diet, 100 μg/kg body weight for mice fed a high-fat diet (Sigma-Aldrich, St Louis, MO, USA) was intraperitoneally injected into male mice that had fasted for 4 h. For glucose tolerance tests, D-glucose (2 g/kg body weight) (Sigma-Aldrich, St Louis, MO, USA) was intraperitoneally injected into male mice that had fasted overnight. For insulin tolerance test, insulin (0.75 U/kg body weight for mice fed a normal chow diet, 1.5 U/kg body weight for mice fed a high-fat diet) (Sigma-Aldrich, St Louis, MO, USA) was intraperitoneally injected into male mice that had fasted for 4 h. For pyruvate tolerance test, pyruvate (2 g/kg body weight) (Sigma-Aldrich, St Louis, MO, USA) was intraperitoneally injected into male mice that had fasted overnight. For epinephrine stimulation test, epinephrine (3µg/10g) (Sigma-Aldrich, St Louis, MO, USA) was intraperitoneally injected into male mice that had fasted for 4h. For immobilization stress, male mice were restrained in a ventilated acrylic restrainer fit to allow the animal to breathe but not to move otherwise.

### Preparation of mouse primary hepatocytes

Primary hepatocytes were isolated from 8-week-old C57BL/6 male mice as previously described (Koo, Flechner et al., 2005). Briefly, mice were anesthetized with avertin (intraperitoneal injection of 250 mg/kg body weight), and livers were perfused with a pre-perfusion buffer (140 mM NaCl, 6 mM KCl, 10 mM HEPES, and 0.08 mg/mL EGTA, pH 7.4) at a rate of 7 mL/min for 5 min, followed by a continuous perfusion with collagenase-containing buffer (66.7 mM NaCl, 6.7 mM KCl, 5 mM HEPES, 0.48 mM CaCl_2_, and 3 g/mL collagenase type IV, pH 7.4) for 8 min. Viable hepatocytes were harvested and purified with a Percoll cushion. Then, hepatocytes were re-suspended in complete growth medium, 199 medium containing 10% FBS, 23 mM HEPES and 10 nM dexamethasone, and seeded on collagen-coated plates at a density of 300,000 cells/mL. After a 4-h attachment period, the medium was replaced with complete growth medium before use in any experiments and changed daily.

### Measurement of cAMP concentrations

Levels of cAMP were quantified using a cAMP ELISA kit (Applied Biosystems, Waltham, MA, USA) according to the manufacturer’s protocol. For mouse primary hepatocytes, cells growing in 6-well plates were lysed with IBMX, and cAMP concentrations were quantified. Concentrations of cAMP in mouse liver samples were measured using an ELISA and normalized to the wet liver weight.

### Measurement of PKA activity

PKA activity was measured using a PKA kinase activity assay kit (Abcam, Cambridge, UK), according to the manufacturer’s protocol. Briefly, whole-cell lysates were incubated with a specific synthetic peptide as a substrate for PKA and a polyclonal antibody that recognizes the phosphorylated form of the substrate. Relative PKA activity was measured by determining the optical density.

### Proximity ligation assay

An *in situ* proximity ligation assay was used to detect protein-protein interactions in cells. Briefly, cells were cultured on collagen-coated confocal dishes and fixed with 4% paraformaldehyde. Then, cells were permeabilized with 0.1% Triton X-100 in phosphate buffer and incubated with blocking buffer for 30 min to prevent nonspecific binding. Samples were sequentially incubated with a primary antibody, PLA probe, ligase, and polymerase, according to the manufacturer’s instructions. PLA-positive cells exhibited red fluorescent signals. The fluorescent signal was observed using a confocal laser scanning microscope (Carl Zeiss 700, 40 X water objectives).

### Quantitative real-time PCR

RNA (2 μg) was reverse transcribed to cDNAs using random hexamer primers, oligo dT and Reverse Transcription Master Premix (ELPIS Biotech, Daejeon, Korea). Quantitative real-time PCR analyses were performed using the cDNAs from the reverse transcription reactions and gene-specific oligonucleotides (Appendix Table S2) in the presence of TOPreal qPCR 2X premix (Enzynomics, Daejeon, Korea). The following PCR conditions were used: an initial denaturation step at 95°C for 10 min, followed by 45 cycles of denaturation at 95°C for 10 s, annealing at 58°C for 15 s and elongation at 72°C for 20 s. The melting curve for each PCR product was assessed for quality control. Supplementary Table 2 shows the sequences of the primers used for qPCR.

### Measurement of hepatic glucose output

Glucose production was determined using the glucose assay kit from Sigma-Aldrich according to the manufacturer’s instructions. Primary hepatocytes were seeded on collagen-coated 6-well plates (0.8 x 10^6^ cells per well). After 3 h, cells were infected with a virus overnight. After 24 h, the medium containing the virus was removed and replaced with fresh 199 medium. After 18 h, the medium was removed, and the cells were rinsed twice with PBS. Then, glucose production buffer (consisting of glucose-free DMEM lacking phenol red, pH 7.4, 20 mM sodium pyruvate, 2 mM l-glutamine and 15 mM HEPES) was added to the cells. After 4 h, the medium was collected and the amount of glucose in the medium was determined using the glucose assay kit (Sigma-Aldrich, St Louis, MO, USA).

### siRNA interference

All siRNAs (Appendix Table S3) were synthesized by Bioneer Inc. (Daejeon, Korea). Cells were transfected with 100 nM siRNA using Lipofectamine RNAiMAX reagent (Invitrogen, Carlsbad, USA).

### Adenovirus preparation and infection

Adenoviruses harboring sh-Con, sh-Rdx (radixin), sh-Prom1, GFP and FERM were produced as previously described (Yi, Park et al., 2013). AD293 cells were re-infected with viral stocks to amplify the viruses, and viruses were purified by double cesium chloride-gradient ultracentrifugation. Infectious viral particles in the cesium chloride gradient (density = ∼1.345) were collected, dialyzed against a 10 mM Tris (pH 8.0), 2 mM MgCl_2_ and 5% sucrose solution and stored at -80°C. Recombinant adenovirus (0.5 × 10^9^ pfu) was injected into the tail veins of mice. Four days after the injection, the mice were subjected to blood glucose metabolism tests.

### Isolation of detergent-resistant lipid rafts

Livers from wild type and Prom1 knockout mice were lysed in 1 mL of lysis buffer (1% Brij 35, 25 mM HEPES, pH 6.5, 150 mM NaCl, protease and protease inhibitor cocktail) and subjected to discontinuous sucrose gradient ultracentrifugation (40%, 30%, and 5%) using a SW41 Ti rotor (39,000 rpm) for 18 h at 4°C. After centrifugation, the sucrose solutions were fractionated into 12 fractions. An opaque buoyant band corresponding to the lipid rafts was collected at the interface between the 30% and 5% sucrose solutions.

### Plasmid construction and transient transfection

Deletion mutants of Prom1-Flag and Myc-radixin (Appendix Table S4) were generated by reverse PCR using the primer sets (Appendix Table S5). mTagBFP2-Farnesyl-5 was a gift from Michael Davidson (Addgene plasmid # 55295). The Myr-BFP-IC3 construct of Prom1 was generated as follows: first, the myristoylation sequence was added at the N-terminus of pmTagBFP2 using reverse PCR; second, Prom1-tail-3xFlag was added at the C-terminus of pmTagBFP2 using the DNA assembly method. Myr-BFP was generated by removing the Prom1-tail from Myr-BFP-IC3 by reverse PCR. DNA transfection was performed using Lipofectamine 3000 reagent (Invitrogen, Carlsbad, USA), according to the manufacturer’s protocol.

### Immunoblotting and immunofluorescence staining

Cells were lysed with the following lysis buffer: 50 mM Tris-Cl, pH 8.0, 150 mM NaCl, 1% NP-40, 0.5% sodium deoxycholate, 0.1% SDS, protease inhibitor mixture and phosphatase inhibitor mixture (Sigma-Aldrich, St Louis, MO, USA). Whole cell lysates obtained from the supernatant after microcentrifugation at 14,000 rpm for 15 min at 4°C were subjected to sodium dodecyl sulfate polyacrylamide gel electrophoresis (SDS-PAGE). The separated proteins were transferred to a nitrocellulose membrane and incubated with specific primary antibodies and horseradish peroxidase (HRP)-conjugated secondary antibodies. Antigens were visualized using an enhanced chemiluminescence substrate kit (Thermo Fisher Scientific, Waltham, MA, USA). For immunoprecipitation, cells were lysed in buffer containing 20 mM Tris-HCl (pH 7.4), 137 mM NaCl, 1 mM MgCl_2_, 1 mM CaCl_2_ and a protease inhibitor cocktail (Sigma-Aldrich, St Louis, MO, USA). Whole-cell lysates (500 μg of protein) were incubated with specific antibodies overnight and then with 60 μL of a slurry of Protein A- or Protein G-agarose beads (Roche, Mannheim, Germany) for 3 h. Immunoprecipitates were analyzed by immunoblotting. For immunofluorescence staining, cells were fixed with PFA and stained with antibodies against the indicated protein.

### Immunohistochemistry

Each paraformaldehyde-fixed samples were either embedded in paraffin or frozen in OCT compound and cut into 5um-thick sections. Tissue samples were then stained with hematoxylin-eosin (HE) or periodic acid-schiff (PAS) according to standard protocol, and the images were captured on light microscope (Leica). For Prom1 immunohistochemistry, fresh-frozen tissues post-fixed in 4% paraformaldehyde. Tissue sections were immunostained with antibody directed against prominin-1 (Thermo Fisher Scientific, Waltham, MA, USA). The samples were pretreated with 2.5% horse serum for 20min to block nonspecific antibody binding and incubated with the antibodies of interest for overnight at 4°C. The slides were then treated with Rhodamine-conjugated secondary antibody. After mounting with fluorescence mounting medium (Agilent, Santa Clara, CA, USA), the samples were captured on an LSM 800 META confocal microscope (Zeiss, Oberkochen, DE).

### Rap activation assay

Rap activity assays were performed as manufacture’s protocol (Abcam; ab212011, Cambridge, UK). Briefly, primary mouse hepatocytes were lysed at 4℃ in a buffer containing 50 mM Tris-Cl, pH 8.0, 150 mM NaCl, 1% NP-40, 0.5% sodium deoxycholate, 0.1% SDS, protease inhibitor mixture and phosphatase inhibitor mixture. Cells lysates were incubated for 30 min with agarose beads coupled to the Rap-binding domain (RBD) of RalGDS (Ral Guanine Nucleotide Dissociation Stimulator), which bind specifically to the active form of Rap. Subsequently, the precipitated GTP-Rap was detected by western blot analysis using anti-Rap1 antibody.

### Statistical analysis

Values are presented as means ± SEM. A two-tailed Student’s t-test was used to calculate the P values

## Acknowledgement

We thank all members of our laboratory for their supports and intellectual inputs during the preparation of this manuscript. Funding: This work was supported by grants from the National Research Foundation of Korea awarded to; Y.-G. Ko (2015R1A5A1009024), H. Lee (2017R1A6A3A01009334) and S. Lee (2017R1D1A1A02017979), and partially by grant from Korea University to H. Lee. Author contributions: Study concept and design, HL, DY, SL and YK; acquisition of data, HL, DY, JP and HYL; analysis and interpretation of data, HL, DY, JL, SL and YK; material support, SK and KS; drafting of the manuscript, SL and YK. YK is the guarantor of this work and, as such, had full access to all the data in the study and takes responsibility for the integrity of the data and the accuracy of the data analysis. Conflicts of interest: The authors disclose no conflict of interests.

## Abbreviations

AKAP: A kinase-anchored protein
AC: adenylyl cyclase
cAMP: cyclic adenosine monophosphate
PKA: protein kinase A
CREB: cAMP response element binding protein
pck: phosphoenol pyruvate carboxykinase
*g6p*: glucose-6-phosphatase
*G6pc*: g6p catalytic subunit
PI3K: phosphatidylinositide 3-kinases
*Fbp1*: fructose 1,6-bisphosphatase 1
GPCR: G protein– coupled receptor
PDE: phosphodiesterase
IBMX: 3-isobutyl-1-methylxanthine
8-Br-cAMP: 8-Bromo-cAMP
DIO: diet-induced obese
shRNA: short hairpin RNA
FERM: F for 4.1 protein, E for ezrin, R for radixin and M for moesin
CFTR: fibrosis transmembrane conductance regulator
NHE3: Na^+^-H^+^ exchanger 3
Cx43: connexin 43
EPAC: exchange protein directrly activated by cAMP

---

**Figure S1.**
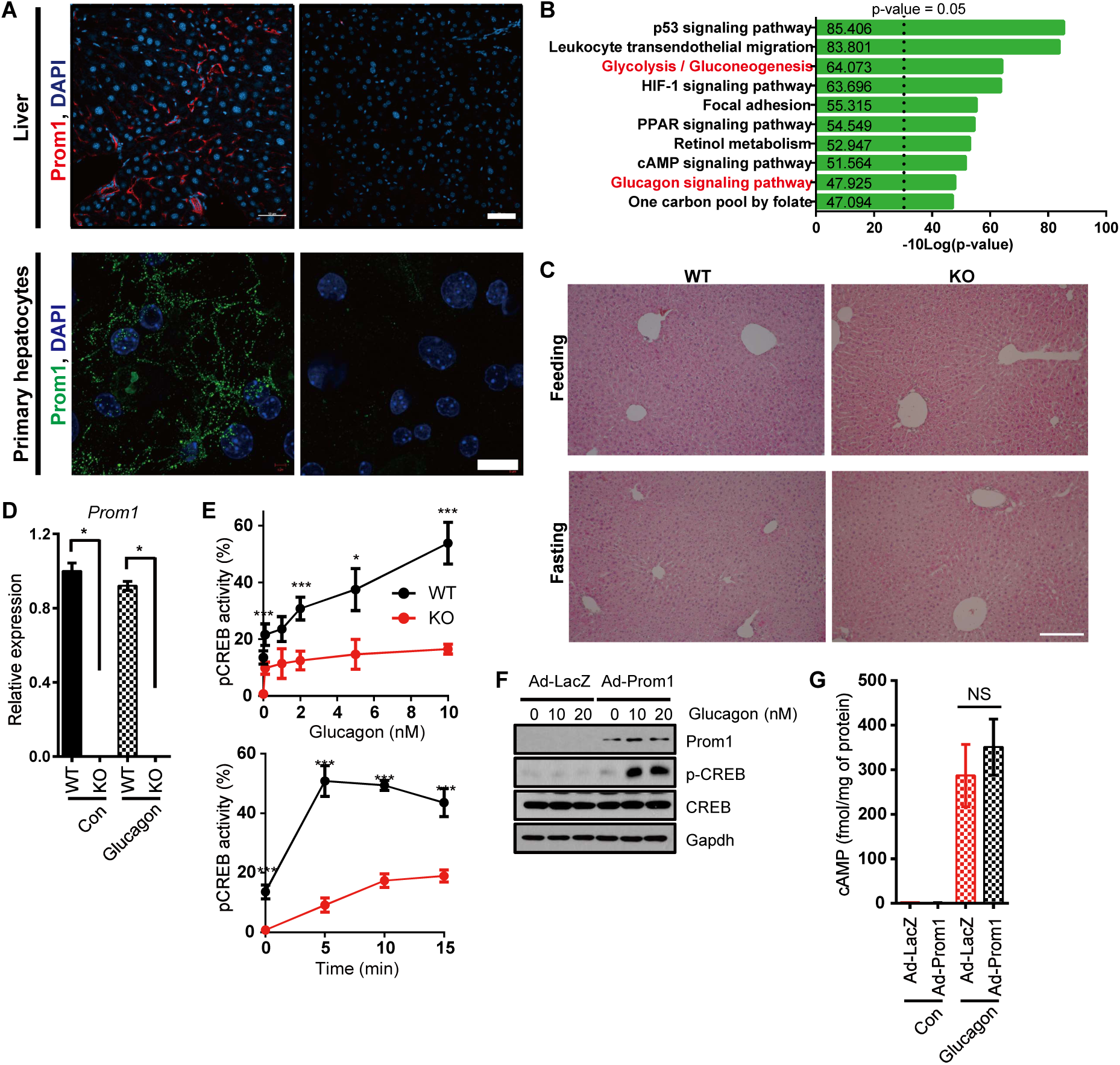
Prom1 disruption prevents glucagon-induced gluconeogenesis in primary mouse hepatocytes. **A** Immunofluorescence staining for Prom1 in the liver and non-permeabilized primary hepatocytes. **B** Enriched KEGG pathway terms for differentially expressed genes. **C** H&E stain (Hematoxylin and eosin stain) of the liver of mouse in each condition. **D** Primary mouse hepatocytes were prepared from 12-week-old male *Prom1*^+/+^ and *Prom1*^-/-^ mice and serum-starved for 16 h. Relative expression levels of *Prom1* mRNAs were determined using RT-PCR. n = 3/group. \*\*\**p* < 0.001. **E** The percentage of cells with nuclear p-CREB staining among more than 300 cells from each group was statistically determined after stimulation with the indicated concentrations of glucagon for 10 min (left panel) or stimulation with 10 nM glucagon for the indicated times (right panel). n = 3/group. \*\*\**p* < 0.001. **F, G** *Prom1* overexpression in *Prom1*^-/-^ hepatocytes restores glucagon-induced CREB phosphorylation. Primary hepatocytes were isolated from 12-week-old male *Prom1*^-/-^ mice, infected with an adenovirus harboring *Prom1* for 24 h, serum-starved for 18 h, and then stimulated with glucagon (10 nM). (**F**) Levels of p-CREB and CREB were determined by immunoblotting 0, 10 and 20 min after glucagon (10 nM) stimulation. (**G**) The cAMP concentration was determined using the cAMP assay kit 10 min after glucagon stimulation (10 nM) in the presence of 10 μM IBMX. n = 3 mice per group. Data information: Data are presented as mean values ± SEM. Two-tailed Student’s t-test was used for statistical analysis. Scale bar; A (top panel, bottom panel) = 50, 20 μm and B = 500 μm.

**Figure S2.**
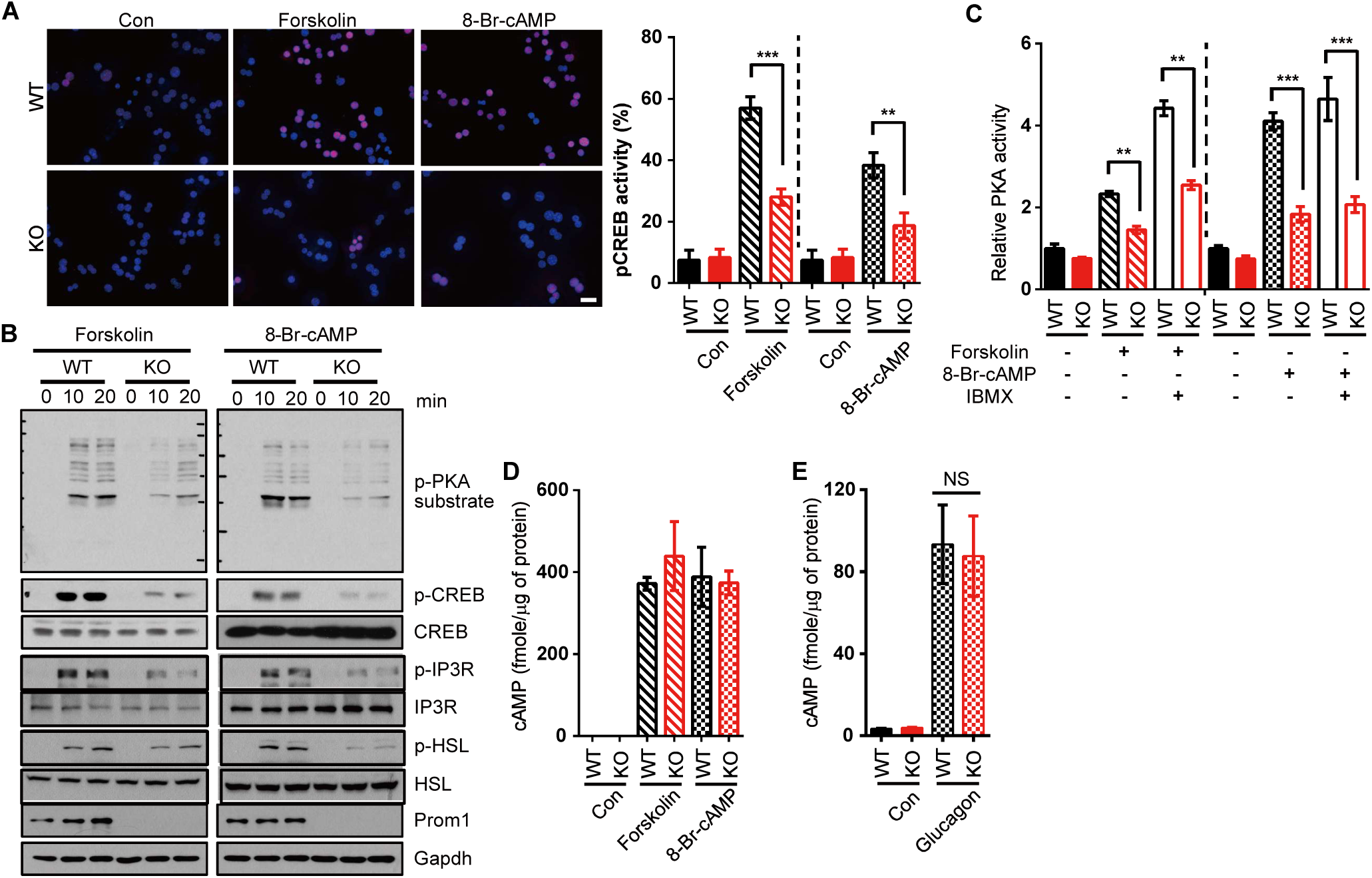
Prom1 disruption prevents glucagon-, forskolin- or 8-Br-cAMP-induced PKA activation in primary mouse hepatocytes. Primary mouse hepatocytes were prepared from 12-week-old male *Prom1*^+/+^ and *Prom1*^-/-^ mice and serum-starved for 16 h. **A** Top panel, the nuclear localization of p-CREB was determined by immunofluorescence staining after a 10-min forskolin (10 μM) or 8-Br-cAMP stimulation (10 μM). Bottom panel, the percentage of cells with nuclear p-CREB staining among more than 300 cells in each group was statistically determined in the images of immunofluorescence staining shown above. \*\**p* < 0.01, \*\*\**p* < 0.001. **B** Levels of p-PKA substrates, p-CREB, CREB, p-IP3R, IP3R, p-HSL, HSL and Prom1 were determined by immunoblotting after stimulation with forskolin (10 μM) or 8-Br-cAMP (10 μM) for the indicated times. **C** Relative PKA activities were determined using the PKA assay kit after 10-min stimulation with 8-Br-cAMP (10 μM) or forskolin (10 μM) in the presence of 10 μM IBMX. n = 3/group. \*\**p* < 0.01, \*\*\**p* < 0.001. **D** The cAMP concentration was determined using the cAMP assay kit after a 10-min stimulation with 8-Br-cAMP (10 μM) or forskolin (10 μM) in the presence of 10 μM IBMX. n = 3/group. **E** The cAMP concentration in the liver was determined using the cAMP assay kit after a 5 min glucagon treatment (2 mg/kg body weight). n = 3/group. Data information: Data are presented as mean values ± SEM. Two-tailed Student’s t-test was used for statistical analysis. Scale bar (A) = 20 μm.

**Figure S3.**
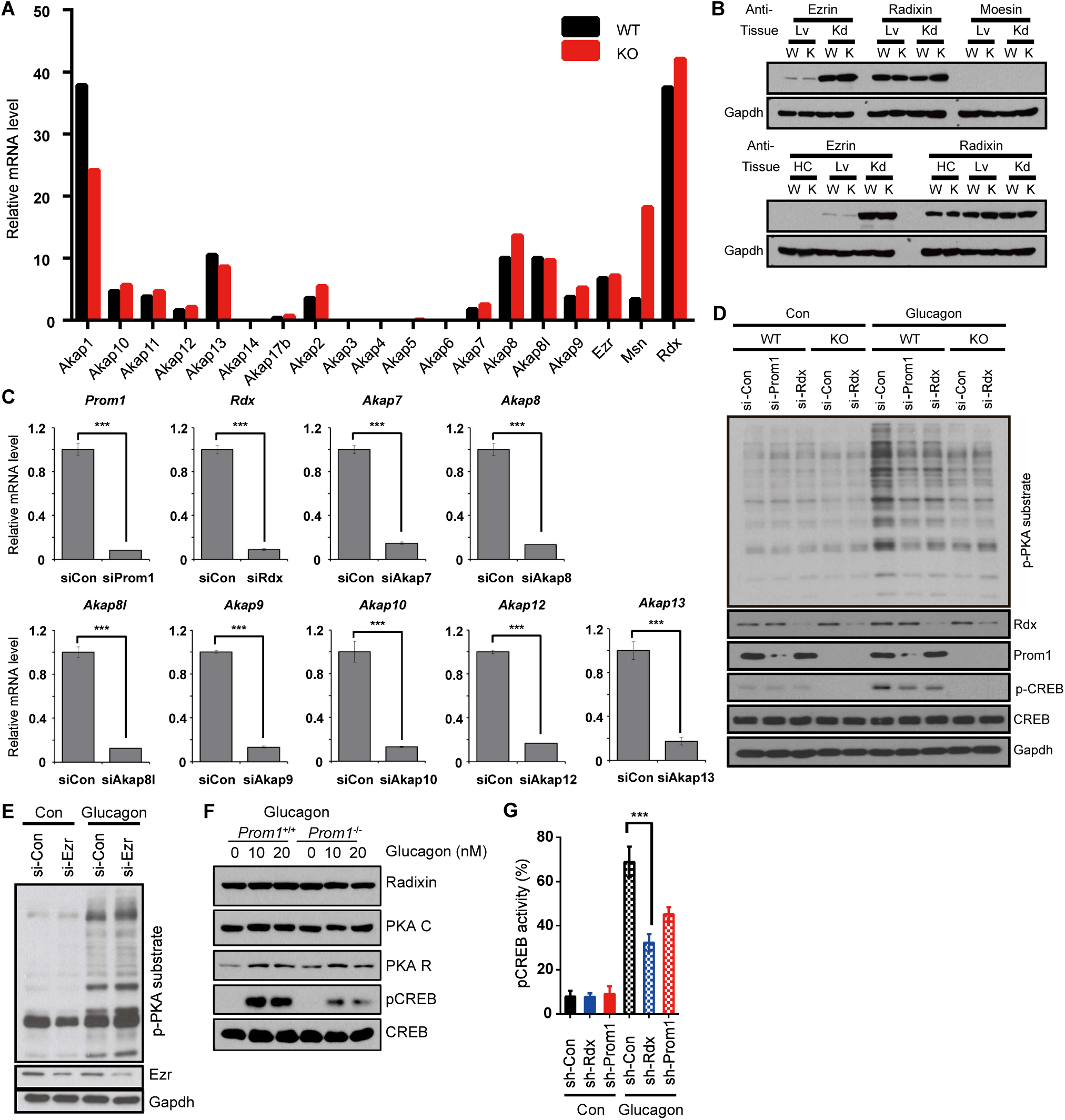
Radixin functions as an AKAP in primary hepatocytes. **A** The relative expression levels of *Akap1, Akap10, Akap11, Akap12, Akap13, Akap14, Akap17b, Akap2, Akap3, Akap4, Akap5, Akap6, Akap7, Akap8, Akap9, ezrin, radixin,* and *moesin* mRNAs in hepatocytes isolated from 12-week-old male Prom1^+/+^ (WT) or Prom1^-/-^ (KO) mice and serum-starved for 16 hours were determined using RNA-seq. AKAP, A-kinase-anchored protein. **B** The expression of ezrin, radixin, and moesin in mouse primary hepatocyte, liver and kidney were determined by immunoblotting. Lv, liver; Kd, kidney; HC, hepatocyte; W, wildtype; K, knockout. **C** The expression of Prom1, radixin, AKAP7, AKAP8, AKAP8I, AKAP9, AKAP10, AKAP12 or AKAP13 was silenced in WT hepatocytes using siRNAs. Hepatocytes were then serum-starved for 18 h and stimulated with glucagon (10 nM) for 10 min. Levels of the *Prom1*, *radixin*, *Akap7*, *Akap8*, *Akap8I*, *Akap9*, *Akap10*, *Akap12* and *Akap13* mRNA were determined using qPCR. n = 3/group. \*\*\**p* < 0.001. **D** Prom1 or radixin expression was silenced in *Prom1*^+/+^ hepatocytes and radixin expression was silenced in *Prom1*^-/-^ hepatocytes. Hepatocytes were serum-starved for 18 h and stimulated with glucagon (10 nM) for 10 min. Levels of p-PKA substrates, p-CREB, CREB, radixin, Prom1 and Gapdh were determined by immunoblotting. **E** Levels of p-PKA substrates, Ezrin and Gapdh in siRNA-treated primary hepatocytes were determined by immunoblotting after stimulation with glucagon (10 nM) for 10 min. **F** Levels of radixin, PKA catalytic subunit, regulatory subunit, p-CREB and CREB were determined by immunoblotting after glucagon stimulation (10 nM). **G** The percentage of cells with nuclear p-CREB staining among more than 300 cells in each group was also statistically analyzed. \*\*\**p* < 0.001. Data information: Data are presented as mean values ± SEM. Two-tailed Student’s t-test was used for statistical analysis

**Figure S4.**
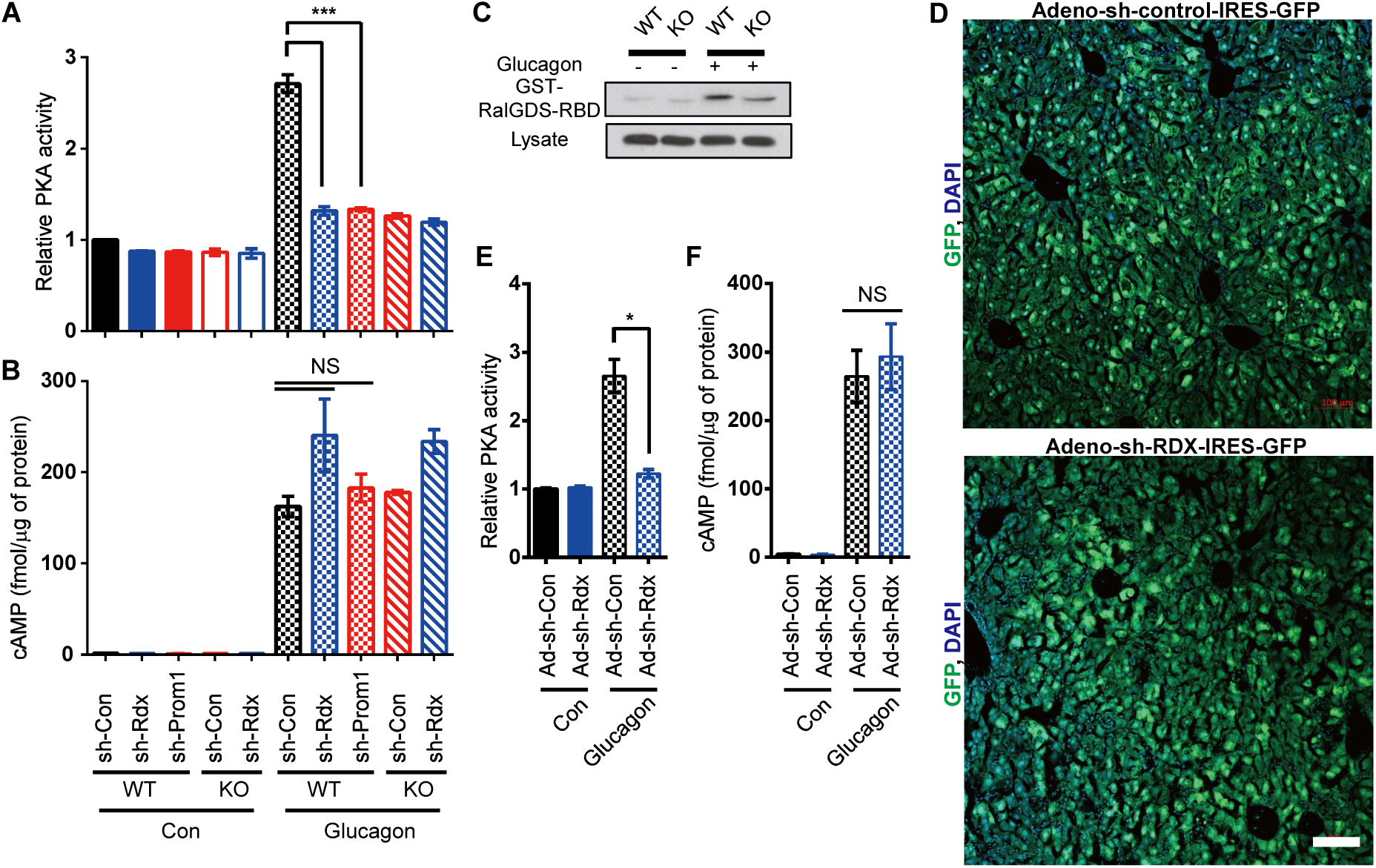
Radixin functions as an AKAP required for glucagon-induced PKA activation in primary hepatocytes and hyperglycemia *in vivo.* **A, B** Prom1 and radixin expression were silenced in *Prom1*^+/+^ (WT) and *Prom1*^-/-^ hepatocytes (KO) by infection with an adenovirus harboring sh-control (sh-Con), sh-radixin (sh-Rdx) or sh-Prom1 for 24 h. Hepatocytes were further serum-starved for 18 h and stimulated with glucagon (10 nM) for 10 min. **A** Relative PKA activities were determined using the PKA assay kit after stimulation with glucagon for 10 min. n = 3/group. \*\*\**p* < 0.001. **B** The cAMP concentration was determined using the cAMP assay kit after a 10-min glucagon stimulation in the presence of 10 μM IBMX. n = 3/group. **C** Pull-down of activated Rap1 with GST-RalGDS-RBD from *Prom1*^+/+^ or *Prom1*^-/-^ primary mouse hepatocytes stimulated with glucagon (10 nM) for 10 min. **D** Confocal microscope images of *Prom1*^-/-^ livers infected with adenovirus harboring sh-Control or sh-Radixin linked with IRES-GFP. **E, F** 12-week-old male wild type mice were infected with an adenovirus harboring sh-control (sh-Con) or sh-radixin (sh-Rdx) for 3 days, fasted for 4 h and intraperitoneally injected with glucagon. **E** Relative PKA activities in the liver were determined using the PKA assay kit 10 min after glucagon (2 mg/kg body weight) stimulation. n = 3/group. \**p* < 0.05. **F** The cAMP concentration in the liver 10 min after glucagon (2 mg/kg body weight) stimulation was determined using the cAMP assay kit. n = 3 per group. Data information: Data are presented as mean values ± SEM. Two-tailed Student’s t-test was used for statistical analysis. Scale bar (D) = 100 μm

**Figure S5.**
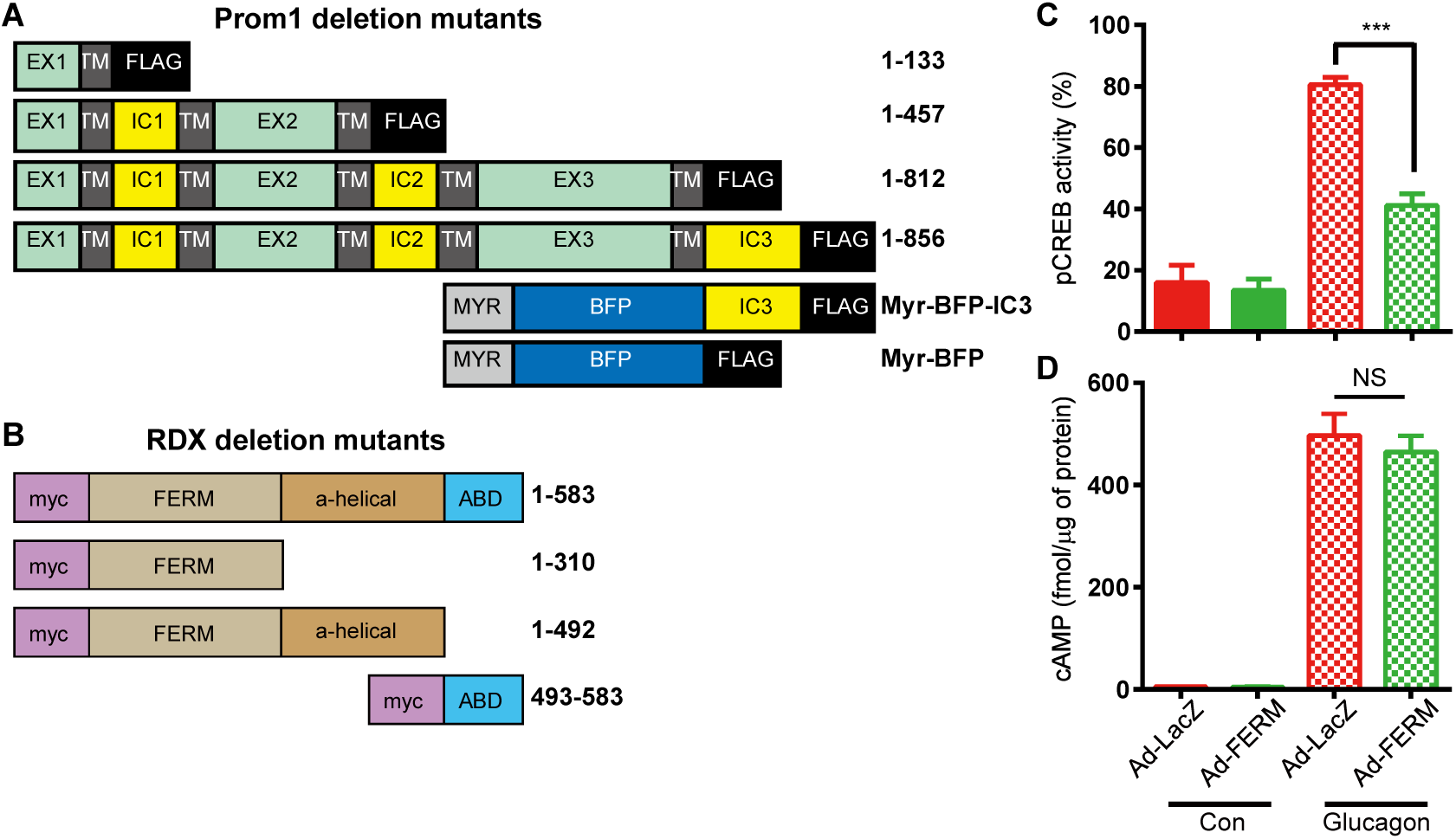
Prom1 and radixin interact via C-terminal cytoplasmic domain of Prom1 and N-terminal FERM domain of radixin. **A, B** Domain structures of deletion mutants of human Prom1 (**A**) and human radixin (**B**). EX, extracellular domain; TM, transmembrane domain; IC, intracellular domain; BFP, blue fluorescence protein; Myr, myristoylation site; FERM, 4.1 protein-ezrin-radixin-moesin domain; ABD, actin-binding domain. **C, D** Overexpression of the FERM domain prevents glucagon-induced PKA activation. Prom1^+/+^ primary hepatocytes were infected with an adenovirus harboring the FERM domain for 48 h, serum-starved for 18 h and stimulated with glucagon (10 nM). **C** The percentage of cells displaying nuclear p-CREB staining among more than 300 cells from each group was statistically determined by analyzing the images of immunofluorescence staining shown in Fig 5E. **D** The cAMP concentration was determined using the cAMP assay kit 10 min after stimulation with glucagon. n = 3 per group. \*\*\**p* < 0.001. Data information: Data are presented as mean values ± SEM. Two-tailed Student’s t-test was used for statistical analysis

